# Follicle stimulating hormone receptor expression in human pancreas and effects on insulin secretion: A translational study

**DOI:** 10.1101/2024.07.12.603223

**Authors:** Banu Kucukemre Aydin, Ceren Incedal Nilsson, Azazul Chowdhury, Quan Wen, Sara Y. Cerenius, Rasmus Stenlid, Katharina Mörwald, Iris Ciba, Hannes Manell, Daniel Weghuber, Anders Forslund, Olof Idevall-Hagren, Peter Bergsten

## Abstract

Follicle-stimulating hormone (FSH) is traditionally known for its role in reproduction, but recent studies suggest it may also influence metabolic processes. This study aimed to examine FSH receptor (FSHR) expression in human pancreatic islets and the direct effects of FSH on insulin secretion, as well as explore FSH’s metabolic role during puberty, focusing on enhanced insulin secretion during this critical period. FSHR gene and protein expression were detected in isolated human pancreatic islets and co-localized with insulin-producing beta-cells. Additionally, FSH at prepubertal (0.1 IU/L) and pubertal concentrations (10 IU/L) significantly enhanced glucose-stimulated insulin secretion (GSIS) and increased intracellular cAMP concentrations in intact human pancreatic islets. In children with obesity from the Beta-JUDO cohort (n=608), plasma FSH levels were positively associated with several insulin secretion indices, particularly in pubertal children. These findings suggest that FSH has significant metabolic roles beyond reproduction, involving insulin secretion and potentially contributing to puberty-related hyperinsulinemia and insulin resistance.

## 1. Introduction

Follicle-stimulating hormone (FSH), synthesized in the anterior pituitary, plays a pivotal role in processes such as puberty, sexual differentiation and reproduction^1^. Additionally, recent studies in perimenopausal women have revealed associations between rising FSH levels and the emergence of various disorders, including obesity, insulin resistance (IR), dyslipidemia, metabolic syndrome (MetS), cardiovascular disease (CVD) and osteoporosis^2–8^. In light of these findings, therapies targeting FSH, such as anti-FSH antibodies have been proposed as a potential future treatment option for menopause related obesity, hypercholesterolemia, and osteoporosis^9^.

Puberty, similar to perimenopause, represents a transitional phase marked by the elevation of gonadotropins, FSH and luteinizing hormone (LH)^10^, along with rapid weight gain^11^, and swift changes in body composition^12^. While an appropriate amount of weight gain is necessary during pubertal development, excessive amounts may increase the risk for obesity, and CVD^13^. The phenomenon of physiological pubertal hyperinsulinemia and IR is well-known^14, 15^. The prevailing view is that hyperinsulinemia and IR play a crucial role in facilitating weight gain and rapid growth in puberty^16^. Although the increase in growth hormone (GH) and insulin-like growth factor 1 (IGF-1) levels during puberty is recognized as an important contributor to this phenomenon^17^, the precise mechanisms and consequences of its alterations are not fully understood^14, 18^.

In a recent study conducted within the Uppsala Longitudinal Study of Childhood Obesity (ULSCO) cohort^19^, we showed that higher plasma FSH levels in prepubertal children with obesity were associated with an increased risk of weight gain and MetS development during puberty. Additionally, we found positive associations between blood insulin and FSH levels during puberty, and hypothesized that rising FSH in this period might have direct effects on insulin secretion via its specific receptor, FSHR, if it is expressed in human pancreatic beta-cells^19^.

FSHR, a G protein coupled receptor with a molecular weight approximately 75 kDa^20^, is primarily expressed on gonads and interacts specifically with FSH^1^. However, previous research revealed extragonadal FSHR expression on mice and human hepatocytes and adipose tissues^21, 22^, and blocking FSH action with specific antibodies reduced body fat in mice^23^. Moreover, FSHR expression and direct regulatory effects of FSH on insulin secretion were reported in rat pancreas^24^, and recently, we presented preliminary results showing FSHR expression in human pancreatic islets and the direct effects of FSH on pancreatic insulin secretion^25^.

In the present study, we further investigated FSHR gene and protein expression in isolated human pancreatic islets, alongside the direct effects of FSH on insulin secretion and FSH-mediated signaling networks. Additionally, we analyzed associations between FSH levels and insulin secretion among children and adolescents in the Beta-cell function in Juvenile Diabetes and Obesity (Beta-JUDO) cohort. Based on previous findings, we hypothesized that FSH may directly influence pancreatic beta-cell function and contribute to the modulation of insulin secretion during puberty, potentially linking FSH to metabolic regulation in this developmental phase.

## 2. Methods

### 2.1 *In vitro* experiments

#### 2.1.1 Isolated human pancreatic islets

Human pancreatic islets were isolated from brain-dead individuals with no known metabolic diseases (Uppsala University Islet Transplantation Unit, Uppsala, Sweden, and Prodo Lab. Inc., Aliso Viejo, CA, USA). Ethical approval for the procedures and protocols regarding the handling of human islets was obtained from the Regional Ethical Review Board in Uppsala, Sweden (EPN number 2010/006; 2010-02-10).

Human pancreatic islets were cultured in Connaught Medical Research Laboratories (CMRL) 1066 medium (Invitrogen, Thermo Fisher Scientific, Waltham, MA, USA) containing 5.6 mM glucose and supplemented with 10% fetal bovine serum (FBS) (Invitrogen, Thermo Fisher Scientific), 100 units/mL penicillin, 100 µg/mL streptomycin and 1% glutamine (Invitrogen, Thermo Fisher Scientific) as previously described^26^. Human pancreatic islets supplied by Prodo Laboratories were cultured in islets specific medium (Prodo Lab. Inc.) for two days before further culture and treatment.

#### 2.1.2 Human embryonic kidney 293 cells (HEK)

HEK cells were used as positive control for FSHR gene- and protein expression since these cells express the receptor^27^. HEK cells were cultured in Gibco Dulbecco’s Modified Eagle Medium (DMEM; Thermo Fisher Scientific), containing 4.5 g/L glucose, supplemented with 10% FBS, 1% Penicillin-Streptomycin and 1% Glutamine.

#### 2.1.3. Real time PCR

Total RNA was extracted from isolated human pancreatic islets and HEK cells by using Macherey-Nagel™ NucleoSpin™ RNA Plus Kit (Thermo Fisher Scientific) according to manufacturer’s instructions. RNA concentration was measured using NanoDrop 2000c Spectrophotometer (Thermo Fisher Scientific). The reverse transcription reaction was performed using iScript™ Reverse Transcription Supermix for RT-qPCR (Bio-Rad Laboratories, Inc. Hercules, CA, US). Obtained cDNA was processed for quantitative real-time reverse transcriptase PCR for FSHR gene (forward primer CTCACCAAGCTTCGAGTCATCCAA and reverse primer AAGGTTGGAGAACACATCTGCCTCT^28^, Invitrogen by Thermo Fisher Scientific). Human Actin (forward primer ACGTGGACATCCGCAAAGAC and reverse primer CAGGGCAGTGATCTCCTTCT) was used as the housekeeping gene. Real time PCR was carried out using Bio-Rad protocol for DNA amplification (Bio-Rad Laboratories) by using IQ™ SYBR Green Supermix (Bio-Rad Laboratories). Products were visualized on a 0.5% agarose gel.

#### 2.1.4. Western blot

Isolated human pancreatic islets and HEK cells were washed and then lysed in Dulbecco’s phosphate-buffered saline (DPBS) (Thermo Fisher Scientific) containing 1% Triton X-100 (Sigma Aldrich, St. Louis, MO, USA). After lysis the samples were centrifuged, and their protein concentration was measured (DC protein assay; Bio-Rad Laboratories). Samples were then electrophoresed and transferred to a PVDF-membrane (Bio-Rad Laboratories) as described previously^29^. Immunoblotting was conducted with anti-FSHR polyclonal antibodies (Host: Rabbit, 1/1000; Invitrogen, Thermo Fisher Scientific).

#### 2.1.5. Immunofluorescence

Isolated human pancreatic islets were hand-selected, embedded in optimal cutting temperature compound (Histolab OCT Cryomount, Histolab, Gothenburg, Sweden), and subsequently frozen. Cryostat microtome (CryoStar NX70, Thermo Fisher Scientific) was used to obtain 5 µm-thick sections. Slides were stored in −80 ^0^C until further analysis. Frozen islet sections were air dried for 30 minutes and then fixed in 2% formaldehyde and sequential 25%, 50%, 75% and 100% methanol. Slides were incubated overnight at 4°C in wet chambers with anti-FSHR polyclonal antibodies (Host: Rabbit, 1:100; Invitrogen, Thermo Fisher Scientific), anti-insulin polyclonal antibodies (Host: Guinea pig, 1:100; Dako, Santa Clara, CA, US) and Alexa Fluor® 488 anti-glucagon antibodies (Host: Mouse, 1:100; Invitrogen, Thermo Fisher Scientific). The second day, after washing, the cells were incubated with Alexa Fluor® 647 anti-rabbit (1/100; Invitrogen, Thermo Fisher Scientific) and Alexa Fluor® 568 anti guinea pig (1/200; Invitrogen, Thermo Fisher Scientific) for 1 hour at room temperature. All antibodies were prepared in 10% bovine serum albumin (BSA) to prevent nonspecific binding. After washing, the slides were counterstained with DAPI (Thermo Fisher Scientific) to stain nuclei and subsequently, the slides were imaged using a Zeiss LSM780 confocal microscope (Carl Zeiss AG, Jena, Germany).

#### 2.1.6. Measurements of glucose-stimulated insulin secretion (GSIS)

Direct effects of FSH on insulin secretion were measured dynamically at normal (5.5 mM) and high (11 mM) glucose concentrations and in the absence or presence of 3 different concentrations of FSH (Sigma-Aldrich); prepubertal FSH concentration (0.1 IU/L), pubertal FSH concentration (10 IU/L) and menopausal FSH concentration (100 IU/L)^6, 30^.

Krebs-Ringer bicarbonate HEPES (KRBH) buffer consisting of 130 mM NaCl, 4.8 mM KCl, 1.2 mM MgSO_4_, 1.2 mM KH_2_PO_4_, 2.5 mM CaCl_2_, 5 mM NaHCO_3_ and 5 mM HEPES was prepared, then titrated to pH 7.4 with NaOH and supplemented with 1 mg/mL BSA and 5.5 or 11 mM glucose.

Approximately 30 human islets were hand-picked for each perifusion chamber and perifused with 5.5 mM glucose containing KRBH buffer at a rate of 130 µL/min, as previously described^26^. After one hour stabilization at 37℃, perifusate samples were collected for 60 minutes. During the first 20 minutes in the presence of 5.5 mM glucose alone, samples were collected with 5 minutes intervals. During the second 20 minutes in the presence of 5.5 mM glucose and in the absence (control) or presence of 3 different doses of human FSH (0.1 IU/L, 10 IU/L and 100 IU/L), samples were collected with 5 minutes intervals. During the last 20 minutes, the glucose concentration was increased to 11 mM with the continued presence of FSH (3 different doses, 0.1 IU/L, 10 IU/L and 100 IU/L) and control condition (one line without FSH), and samples were collected more frequently (2^nd^, 4^th^, 6^th^, 10^th^, 15^th^, and 20^th^ minutes). After perifusion, islets were rinsed with DPBS and lysed in DPBS containing 1% Triton X-100. Perifusate samples and lysed islets were stored at −20℃ until further analysis.

Ultrasensitive human insulin ELISA kits were used for the measurement of insulin levels (Mercodia AB, Uppsala, Sweden). A standard curve was constructed from the insulin standards provided in the kit. The absorbance of the 96-well ELISA culture plate was read in a Multiscan Go plate reader (Thermo Fisher Scientific).

Total protein content in the lysates was determined by detergent compatible protein assay (DC protein assay, Bio-Rad Laboratories), according to the Lowry method^31^. Insulin concentration of the perifusate samples was normalized by using total protein content and insulin concentrations in the lysates.

#### 2.1.7. Investigation of FSH-mediated signaling networks in human pancreatic islets

FSHR is a G-protein coupled receptor, and the stimulatory Gαs protein is a well-known transducer that primarily leads to an increase in intracellular cyclic adenosine monophosphate (cAMP)^32^. Accordingly, to test whether FSH stimulates cAMP formation in human pancreatic islet cells, we transduced the islets with adenoviral vectors encoding the cAMP sensor EpacS^H187^ and the Ca^2+^ sensor R-GECO1, both under the control of a general CMV promoter enabling expression in all islet cell types^33, 34^.

The human islets were handpicked under stereomicroscope and cultured in CMRL medium with 5.5 mM glucose supplemented with 10% FBS, 100 U/mL penicillin and 100 μg/mL streptomycin at 37°C and 5% CO_2_. For expression of the Ca^2+^ sensor R-GECO1 and the cAMP biosensor EpacS^H187^, 15-30 islets were infected with 2.5 µL high titration virus (>10^12^–10^13^ vp/mL; Vector Biolabs Malvern, PA) in 200 µL of culture medium and incubated for 3 hours at 37°C in a humidified atmosphere containing 5% CO_2_. After incubation, 3 mL of culture medium was added, and the islets were incubated for 24-48 hours to allow for expression of the fusion protein. On the day of experiment, islets were handpicked and placed on a poly-L-lysine-coated coverslip mounted in an open chamber and perifused with a buffer containing 125 mM NaCl, 4.9 mM KCl, 1.2 mM MgCl_2_, 1.3 mM CaCl_2_, 25 mM Hepes, 5.5 mM glucose, and 0.1% BSA at a pH of 7.4. The chamber was placed on the stage of a Nikon TiE microscope equipped with temperature control to keep the islets at 37°C. Imaging was performed using a TIRF illuminator and a 60×/1.45 NA plan-Apo objective (Nikon, Tokyo, Japan). The system utilized diode-pumped solid-state lasers (Cobolt, Hübner photonics, Solna, Sweden) to provide excitation light for mTurquoise2 and R-GECO (445 nm and 561 nm, respectively) and emission light was separated by interference filters (483/32 for mTq2, 542/27 for mCitrine, 597LP for R-GECO1). cAMP concentration changes were determined by calculating the 483/542 FRET ratio. Image analysis was performed offline using ImageJ. Briefly, islet cells were manually identified, and regions of interest were drawn within the footprints of individual islet cells. The fluorescence intensity changes for donor (483 nm emission), acceptor (542 nm emission) and R-GECO (597 nm) was measured over time and values were normalized to the pre-stimulatory value (5.5 mM glucose). Only cells with a robust response to 10 IU/L FSH were included in the analysis (55 % of all cells).

#### 2.1.8. Statistical analyses of the islet study

Comparisons between the groups were done by using one-way ANOVA. Dunnett’s test was used for post hoc analyses. Results are presented as mean ± SEM, unless otherwise stated. Analyses were performed by using GraphPad Prism (GraphPad Software Inc., La Jolla, CA). Statistical significance was defined as p< 0.05.

### 2.2. The Beta-JUDO cohort study

#### 2.2.1. Subjects, design and setting

Beta-JUDO is an ongoing, dual-center longitudinal cohort study with children and adolescents from the Uppsala University Hospital (ULSCO cohort, Uppsala, Sweden), and the Paracelsus Medical University Hospital cohort (Salzburg, Austria). The cohort was designed to understand and define factors contributing to childhood obesity, development of obesity-related metabolic diseases as well as the effects of interventions and includes children and adolescents with overweight/obesity and normal weight^35, 36^. Children with obesity were screened for inclusion in this study during clinical baseline visits at the respective hospital in Uppsala and Salzburg. Normal weight children were enrolled through collaboration with local schools and advertisements^35^.

In the present study, only prepubertal and pubertal children of the Beta-JUDO cohort were included. Postpubertal individuals (n=222) and children with unknown pubertal status (n=14) were excluded from the study. Additionally, children and adolescents who had syndromic obesity, type 1 or type 2 diabetes mellitus, and those who were taking medications which may affect glucose metabolism or gonadotropin levels were not included in the analyses (n=23). An additional six patients were excluded due to missing all relevant anthropometric data and blood measurements. The final study population comprised 608 children and adolescents (Uppsala n=336, Salzburg n=272), aged between 5 to 18 years, including 522 with obesity and 86 without. Of these participants, 185 children were prepubertal at their initial visit, while 423 were in puberty. We further analyzed longitudinal data from the prepubertal children who had follow-up visits during puberty (n=62). However, only seven prepubertal children without obesity attended follow-up visits during puberty, and none underwent an OGTT at these follow-up visits, which prevented longitudinal data analyses for this subgroup. Thus, the final population for longitudinal analyses consisted of 55 children. If medications were required during the follow-up, we used data from the visit prior to the start of treatment. All procedures adhere to the standardized operating procedures, which were harmonized across study centers.

#### 2.2.2. Ethics

The study was performed in accordance with the Declaration of Helsinki. The Beta-JUDO study was approved by the Uppsala Regional Ethics Committee (2010/036 and 2012/318) and the Salzburg Ethics Committee (EK1544/2012). The children and guardians received oral and written information regarding participation in the project. Written consent was obtained from legal guardians. Additionally, adolescents aged 12 years and above provided their own consent. Collected data was stored and retrieved using Research Electronic Data Capture (REDCap) tool^35^, in which the subjects were coded.

#### 2.2.3. Physical examinations and measurements

Weight in kilograms was measured while the patient wore lightweight clothing, employing a standardized, calibrated SECA model 704 scale in Uppsala and model 801 scale in Salzburg (SECA, Hamburg, Germany). Height in centimeters was recorded twice using an Ulmer stadiometer (Busse, Elchingen, Germany) in Uppsala and a SECA model 222 stadiometer (SECA, Hamburg, Germany) in Salzburg, with the average result noted. BMI was then computed as the weight in kilograms divided by the square of height in meters. BMI-SDS was calculated with Microsoft Excel add-in LMS Growth version 2.76 using World Health Organization 2007 growth reference^37^. Waist circumference was measured to the nearest 0.1 cm using a flexible tape. Measurements were taken in the horizontal plane, midway between the inferior costal margin and the iliac crest, with the subject standing and feet placed together. Hip circumference was also measured to the nearest 0.1 cm at the point of maximum girth around the buttocks. Children were evaluated for puberty according to the Tanner classification^38, 39^. Pubertal staging was supplemented through self-description and basal LH levels due to a high refusal rate for pubertal examination. Pubertal onset was determined based on a basal LH level of ≥0.3 IU/L, consistent with previous research^40, 41^.

#### 2.2.4. Blood sampling and measurements

All fasting blood samples were collected between 8:00 and 10:00 am after 10-hour overnight fast and analyzed locally at the respective hospital in Uppsala and Salzburg. OGTT was performed as previously described^35^. Blood was drawn at fasting and at minutes 5, 10, 15, 30, 60, 90, 120, 180 post-ingestion for insulin and glucose measurements. All other parameters were measured only in fasting blood samples. Validation of analyses was performed between the laboratories by using reference blood samples. A correction factor generated from a previous study within the Beta-JUDO cohort was applied to all glucose values to compensate for the use of different blood collection tubes at the two sites^36^. The original Salzburg values were corrected using the following formula [(1.0153 x original Salzburg glucose value in mM) + 0.2489]^36^.

#### 2.2.5. Calculation of insulin secretion and sensitivity measurements

Hyperglycemic and euglycemic-hyperinsulinemic clamp studies are well-established methods for assessing beta-cell function and insulin sensitivity^40^. However, these procedures are invasive, complex, and costly, making them impractical for large-scale use, particularly in children. To calculate insulin secretion and sensitivity, we employed surrogate indices using blood glucose and insulin concentrations measured during the OGTT.

Delta insulin was calculated from OGTT by the formula: insulin_30 min_ – insulin_0 min_. The insulinogenic index (IGI) was calculated as (insulin_30 min_ – insulin_0 min_)/(glucose_30 min_ – glucose_0 min_)^42^. Stumvoll et al.^43^ demonstrated that insulin secretory indices from the OGTT strongly correlate with the first and second phases of insulin release (S1PhOGTT and S2PhOGTT, respectively) as measured by the hyperglycemic clamp. Thus, we applied the formulas to calculate S1PhOGTT (1283 + 1.829 × insulin_30 min_ − 138.7 × glucose_30 min_ + 3.772 × insulin_0 min_) and S2PhOGTT (287 + 0.4164 × insulin_30 min_ − 26.07 × glucose_30 min_ + 0.9226 × insulin_0 min_) to assess first and second phases of insulin release^43, 44^. To evaluate insulin resistance, the homeostatic model assessment for insulin resistance (HOMA-IR) was calculated using the formula: fasting glucose (mmol/L) × fasting insulin (μIU/mL) / 22.5^45^.

#### 2.2.6. Statistical analyses of the Beta-JUDO cohort study

The Shapiro-Wilk test was used to assess the normality of the variable distributions. Given that most variables did not follow a normal distribution, comparisons were made using the Mann-Whitney U test for independent samples or the Wilcoxon Signed Ranks test for paired samples. Variables were presented as medians and interquartile ranges (IQR), and Spearman’s rank correlation coefficients were employed for correlation analyses. The chi-squared test was applied to categorical variables. Separate evaluations for children with and without obesity were performed for comparison and correlational analyses. Multiple linear regression models were constructed to investigate the independent associations of blood FSH levels with insulin secretory indices derived from OGTT results, adjusting for potential confounders. For data not meeting normality criteria, regression analyses were executed on log-transformed variables. All statistical analyses were carried out using SPSS version 15.0 (SPSS Inc., Chicago, IL). A p-value of less than 0.05 was considered statistically significant.

## 3. Results

### 3.1. Human pancreatic islet beta-cells express FSHR

RT-PCR analysis revealed the presence of the FSHR gene product in human islets from three donors, and Western blot analysis confirmed the expression of FSHR at the protein level (Fig. 1). Additional investigation with confocal immunofluorescence microscopy localized FSHR expression to insulin-producing beta-cells in isolated human pancreatic islets (Fig. 2).

**Figure 1.**
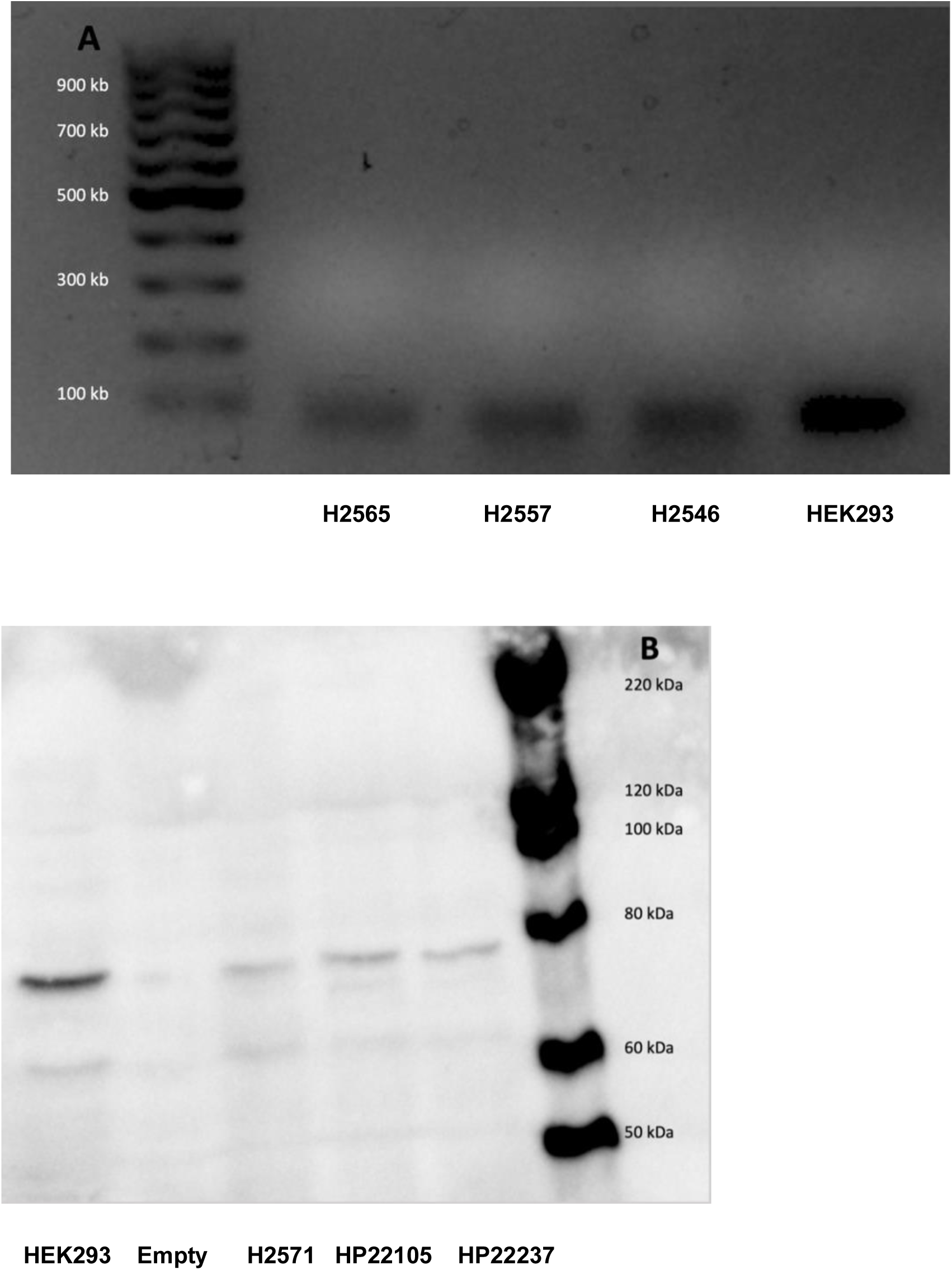
Human pancreatic islets express follicle stimulating hormone receptor (FSHR). FSHR transcript (Panel A) and FSHR protein (Panel B) detected in islets from three donors and HEK293 cells.

**Figure 2.**
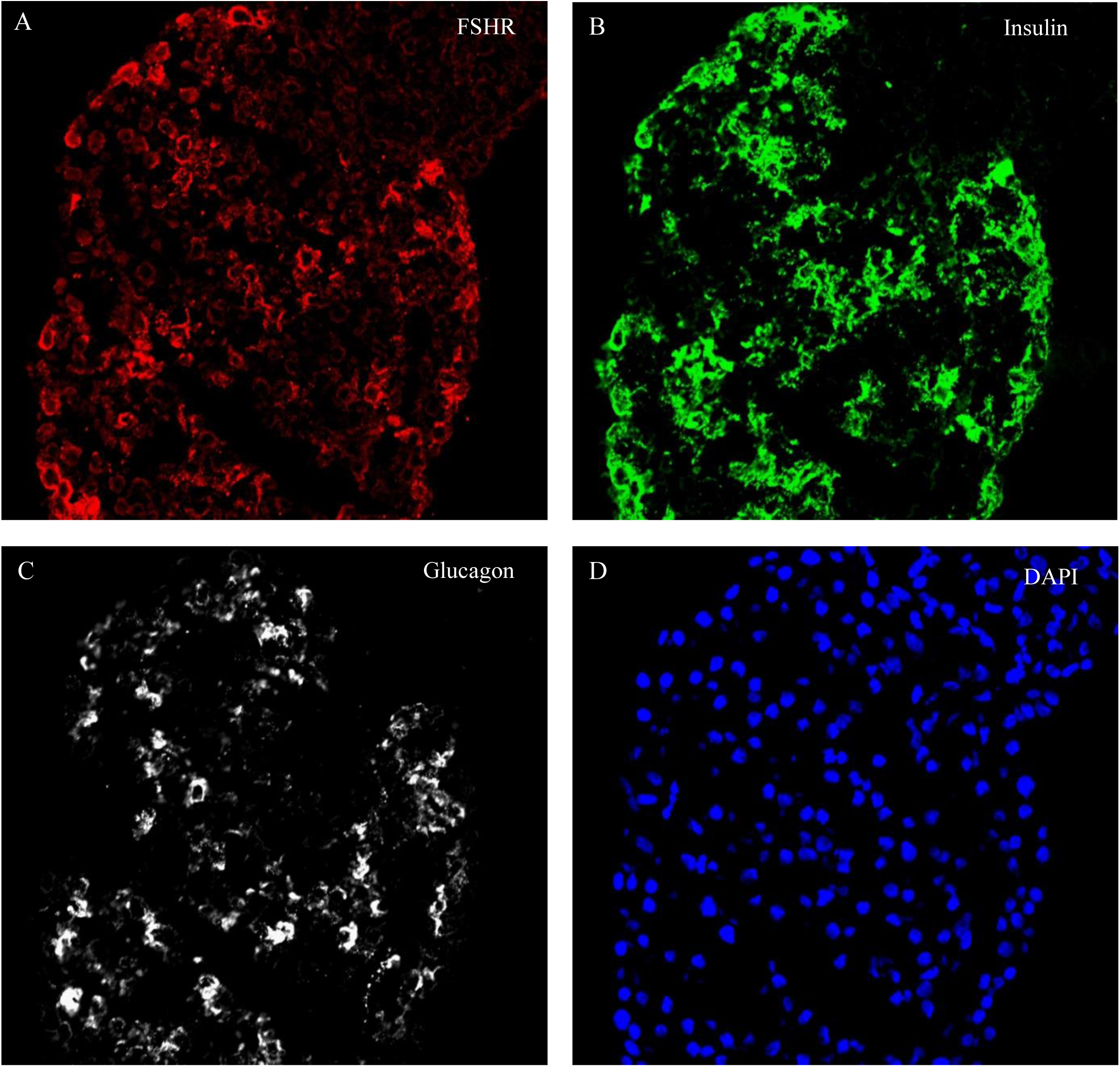
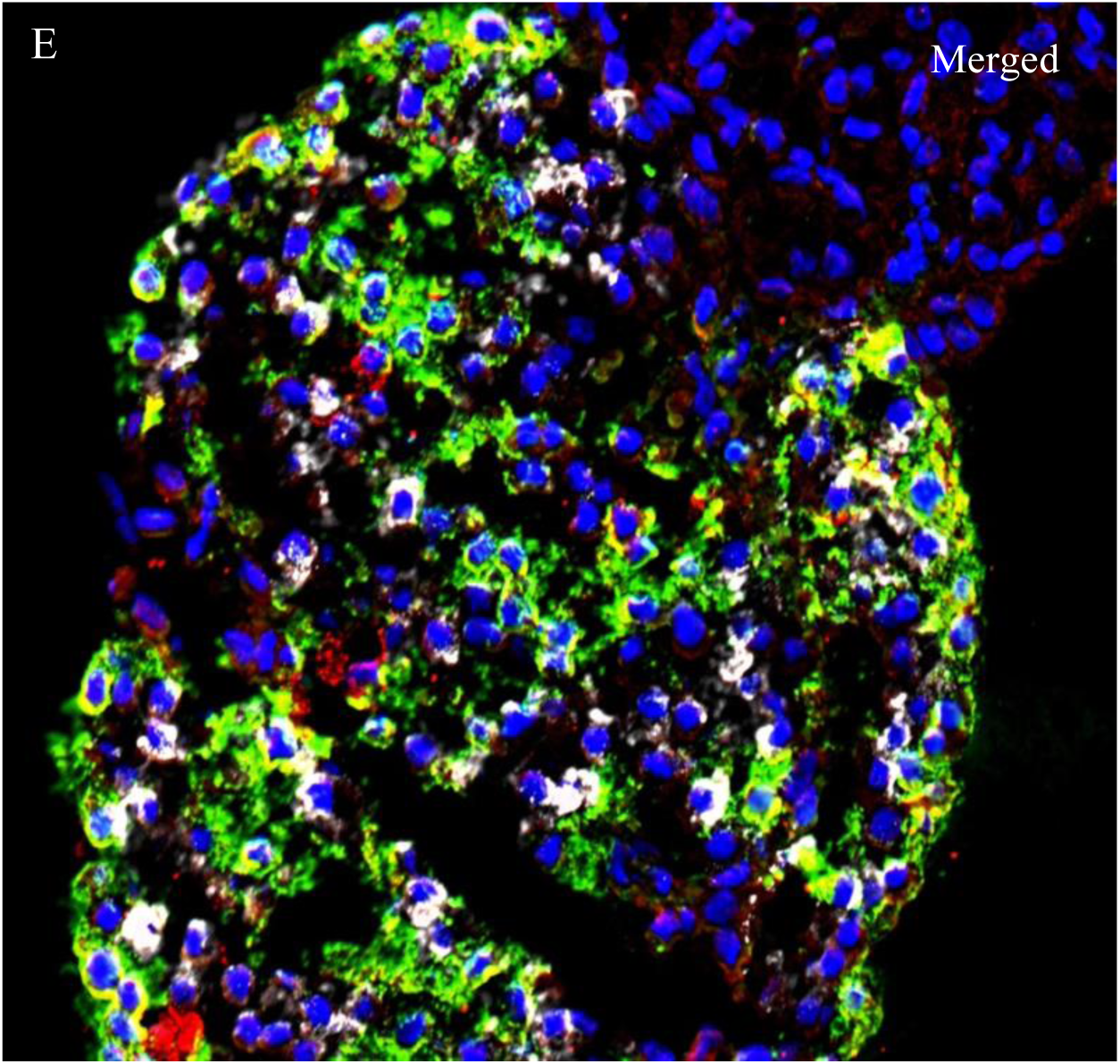
Human pancreatic islet beta-cells express follicle stimulating hormone receptor (FSHR). Human islet sectioned and stained for FSHR (red) (A), insulin (green) (B), glucagon (white) (C) and nuclei/DAPI (blue) (D). Merged picture of FSHR, insulin, glucagon and DAPI (E) is shown. Representative of experiments with five different donors.

### 3.2. FSH accentuated GSIS in isolated human pancreatic islets

We assessed the effect of FSH on insulin secretion under varying glucose concentrations. At a basal glucose concentration of 5.5 mM, the addition of FSH at concentrations of 0.1, 10, or 100 IU/L did not significantly affect insulin secretion (Fig. 3). However, substantial enhancement of insulin secretion was observed at elevated glucose concentration of 11 mM. Particularly, FSH doses of 0.1 IU/L and 10 IU/L amplified GSIS significantly (Fig. 3).

**Figure 3.**
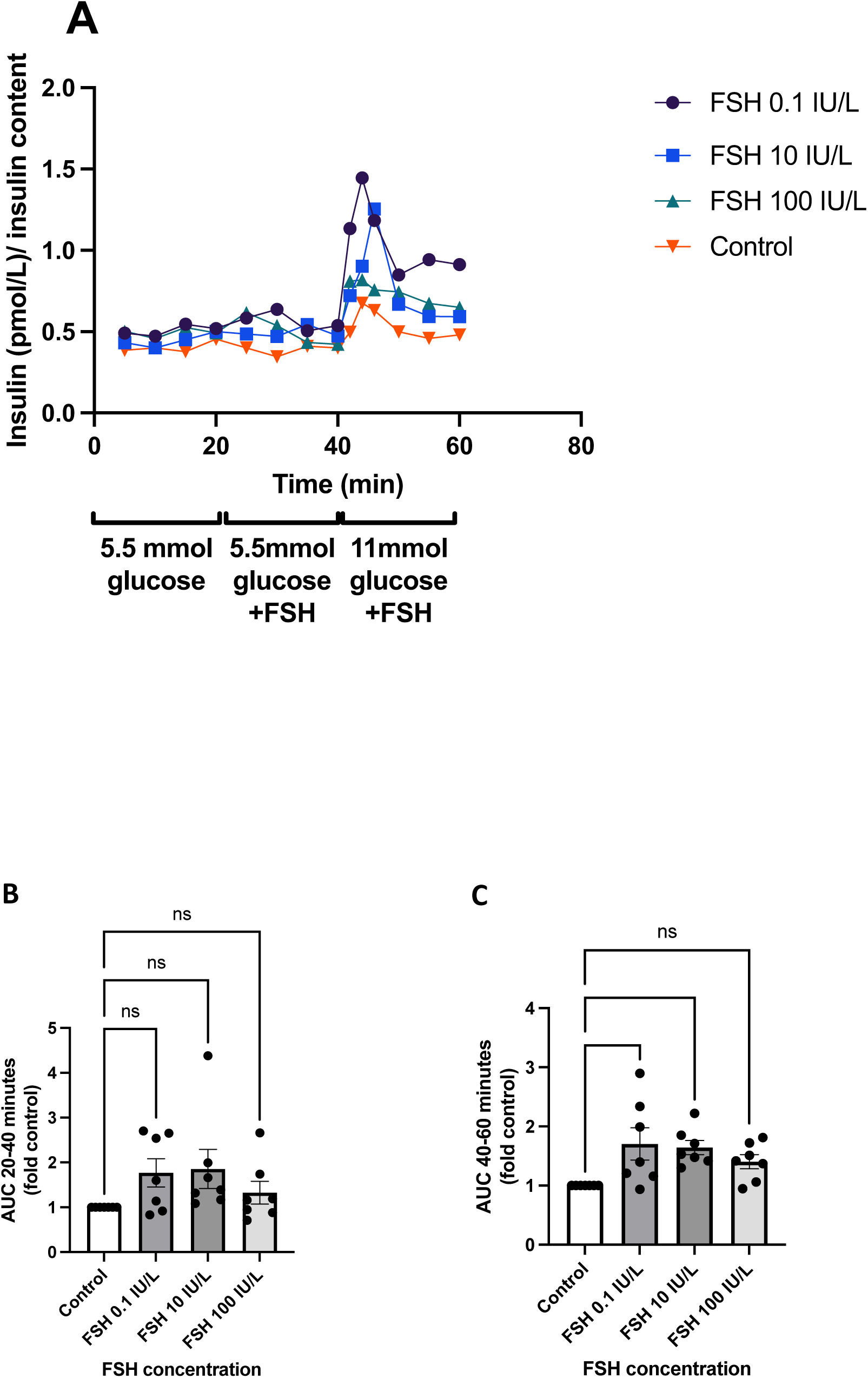
Follicle stimulating hormone (FSH) accentuates glucose-stimulated insulin secretion in isolated human pancreatic islets. Measurement of insulin secretion from isolated human pancreatic islets at 5.5- and 11-mM glucose concentrations in the presence of 0.1, 10 or 100 IU/L FSH and control islets without FSH (Panel A). Comparison of the area under the curve between 20-40 minutes (5.5 mM glucose with different FSH concentrations) (Panel B). Comparison of the area under the curve between 40-60 minutes (11 mM glucose with 3 different FSH concentrations) (Panel C). Results are presented as means ± SEM of seven separate experiments (ns=nonsignificant; *, p=0.01 and p=0.02)

### 3.3. FSH dose-dependently stimulated cAMP formation in isolated human pancreatic islets

Although FSH addition at a basal glucose concentration of 5.5 mM did not significantly change insulin secretion, exposure to 10 IU/L FSH under the same conditions resulted in a distinct increase in the EpacS^H187^ FRET ratio in about 50% of the cells (Fig. 4), indicating a rise in cAMP levels. This response was enhanced in the presence of elevated glucose (11 mM) (Fig. 4). The non-responding cells (Fig. 4) may reflect the other islet cell types, which do not express FSHR. Elevation of the glucose concentration to 11 mM triggered a small increase in cAMP concentration by itself (Fig. 4), as previously shown^46^, and was accompanied by an elevation of the intracellular Ca^2+^ concentration (Fig. 4). In the presence of 11 mM glucose, FSH dose-dependently (0.1 to 100 IU/L) increased cytosolic cAMP concentration which also coincided with enhanced Ca^2+^ responses in the cells (Fig. 4). These results show that activation of FSHR on human islet β-cells results in dose-dependent formation of cAMP.

**Figure 4.**
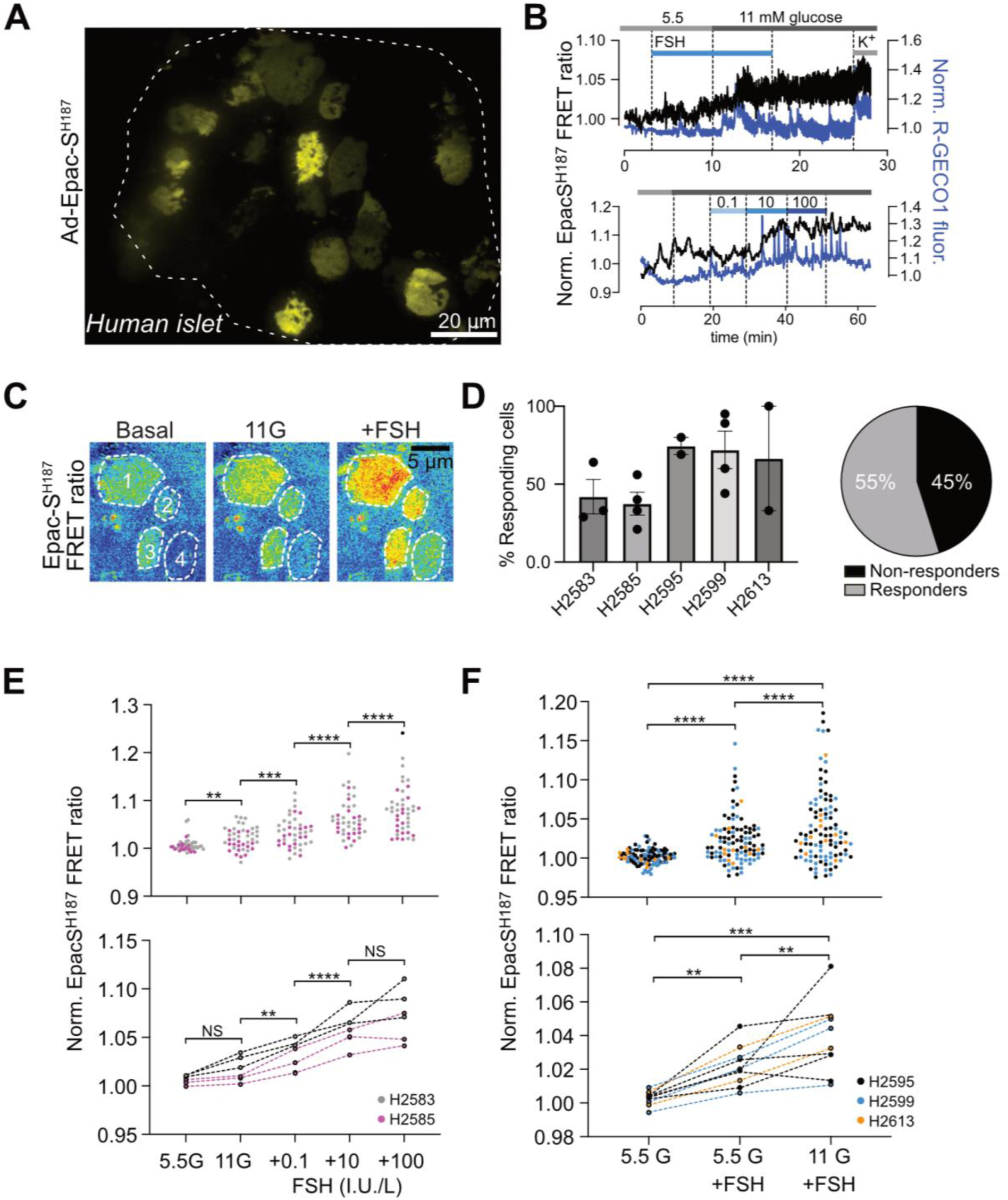
Follicle stimulating hormone (FSH) stimulates cAMP production in a glucose-dependent manner. A. TIRF microscopy image of a human islet expressing the cAMP sensor EpacS^H187^. B. Example recordings of cAMP (black) and Ca^2+^ (blue) from human islet β-cells exposed to the indicated stimuli. C. TIRF microscopy images of part of a human islet showing four cells (outlined by dashed lines) expressing EpacS^H187^ following exposure to 5.5 mM glucose (basal), 11 mM glucose (11G) and 11 mM glucose and 10 IU/L FSH (+FSH). D. Quantification of the number of cells that show an increase in cAMP concentration in response to 10 IU/L FSH. Categories on the x-axis represent human islet donors (N=5 donors; n=15 islets; n=248 cells). E. Normalized EpacS^H187^ FRET ratio changes in human islet cells exposed to 5.5 mM glucose (5.5G), 11 mM glucose (11G) and 11 mM glucose together with FSH at the indicated concentration (0.1, 10, 100 IU/L). Values were extracted after 10 min exposure to the indicated stimuli. Only cells that showed a cAMP increase to 10 IU/L FSH was included in the analysis (see panel C). The upper graph shows individual cells and the lower graph show individual islets (N=2 donors; n=6 islets; n=43 cells). F. Normalized EpacS^H187^ FRET ratio changes in human islet cells exposed to 5.5 mM glucose (5.5G), 5.5 mM glucose and 10 IU/L FSH (5.5G+FSH) and 11 mM glucose and 10 IU/L FSH (11G+FSH). Values were extracted after 10 min exposure to the indicated stimuli. Only cells that showed a cAMP increase to 10 IU/L FSH were included in the analysis (see panel C). The upper graph shows individual cells and the lower graph show individual islets (N=3 donors; n=10 islets; n=105 cells). ** P<0.01; ***P<0.001; ****P<0.0001; 2-way ANOVA with Tukey’s post hoc test.

### 3.4. FSH is positively associated with insulin responses during OGTT

#### 3.4.1. First visit, cross-sectional data analysis of children from the Beta-JUDO cohort

##### 3.4.1.1. Comparative analyses in children without obesity

Table 1 summarizes the clinical and laboratory characteristics of children without obesity at the first visit, categorized by pubertal status. While fasting insulin and HOMA-IR levels were higher in pubertal children, no significant differences were observed in delta insulin, IGI, or the first and second phase insulin responses during the OGTT (S1PhOGTT and S2PhOGTT).

**Table 1.** Cross-sectional comparison of clinical and laboratory characteristics in prepubertal and pubertal children without obesity at their initial visit.

|  | Prepubertal<br>(n=20) | Pubertal<br>(n=66) | p-value* |
| --- | --- | --- | --- |
| Age (years) | 8.5 (7.2-10.0) | 15.0 (13.3-15.6) | **<0.001** |
| Sex (%) | 13 boys (65%)/7 girls (35%) | 42 boys (64%)/ 24 girls (36%) | 0.91 |
| BMI-SDS | -0.2 (-1.0-0.3) | -0.1 (-0.7-0.5) | 0.21 |
| WC/height | 0.44 (0.41-0.46) | 0.43 (0.41-0.44) | 0.23 |
| WC/HC | 0.86 (0.84-0.91) | 0.84 (0.79-0.87) | **0.02** |
| LH (IU/L) | 0.1 (0.1-0.1) | 2.65 (1.68-4.13) | **<0.001** |
| FSH (IU/L) | 0.85 (0.54-1.9) | 3.65 (2.3-5.5) | **<0.001** |
| Estradiol (pmol/L) | 40 (40-40) | 102 (47.8-156.6) | **<0.001** |
| Testosterone | 0.42 (0.42-0.42) | 7.0 (0.99-17.1) | **<0.001** |
| Fasting glucose (mmol/L) | 5.2 (5.0-5.4) | 5.0 (4.7-5.4) | 0.11 |
| OGTT glucose 5 min (mmol/L) | 5.5 (5.1-6.0) | 5.4 (4.9-6.0) | 0.32 |
| OGTT glucose 10 min (mmol/L) | 6.6 (6.2-7.2) | 6.2 (5.6-6.9) | 0.29 |
| OGTT glucose 15 min (mmol/L) | 7.4 (6.0-7.8) | 7.18(6.38-7.87) | 0.99 |
| OGTT glucose 30 min (mmol/L) | 7.7 (7.3-8.5) | 8.0 (7.4-9.2) | 0.48 |
| OGTT glucose 60 min (mmol/L) | 7.0 (5.8-8.2) | 6.6 (5.8-7.5) | 0.33 |
| OGTT glucose 90 min (mmol/L) | 6.5 (6.3-7.0) | 6.1 (5.6-6.7) | **0.01** |
| OGTT glucose 120 min (mmol/L) | 6.6 (5.9-7.5) | 6.2 (5.6-6.9) | 0.18 |
| Fasting insulin | 33 (20-54) | 49 (41-61) | **0.007** |
| OGTT insulin 5 min | 115 (33-195) | 77 (56-137) | 0.88 |
| OGTT insulin 10 min | 201 (51-326) | 201 (118-271) | 0.84 |
| OGTT insulin 15 min | 278 (77-347) | 288 (181-392) | 0.37 |
| OGTT insulin 30 min | 320 (153-469) | 368 (266-550) | 0.28 |
| OGTT insulin 60 min | 299 (220-328) | 291 (188-451) | 0.59 |
| OGTT insulin 90 min | 243 (208-335) | 236 (146-292) | 0.58 |
| OGTT insulin 120 min | 181 (107-351) | 208 (139-292) | 0.59 |
| Delta insulin 0-30 min | 260 (123-411) | 324 (214-494) | 0.25 |
| Insulinogenic index | 0.96 (0.40-1.35) | 0.74 (0.56-1.35) | 0.87 |
| S1PhOGTT | 1050 (607-1222) | 1033 (802-1330) | 0.54 |
| S2PhOGTT | 279 (177-322) | 278 (223-343) | 0.52 |
| HOMA-IR | 1.03 (0.60-1.81) | 1.58 (1.09-2.01) | **0.03** |

Abbreviations: BMI-SDS: Body mass index standard deviation score; FSH: Follicle-stimulating hormone; HC: Hip circumference; HOMA-IR: Homeostatic model assessment for insulin resistance; LH: Luteinizing hormone; min: Minutes; OGTT: Oral glucose tolerance test; S1PhOGTT: Stumvoll index 1; S1PhOGTT: Stumvoll index 2; WC: Waist circumference. All values are median and (Q1-Q3 interquartile ranges), except if otherwise stated. \*Bold text indicates a statistically significant difference with a p-value less than 0.05.

##### 3.4.1.2. Correlation analyses in children without obesity

Plasma FSH concentrations were positively correlated with delta insulin (r=0.308, p=0.02), IGI (r=0.277, p=0.04), S1PhOGTT (r=0.339, p=0.009), S2PhOGTT (r=0.344, p=0.008) and HOMA-IR (r=0.319, p=0.006). Additionally, FSH levels showed positive correlations with fasting insulin (r=0.376, p=0.001), and insulin values during the OGTT at 15 minutes (r=0.300, p=0.02) and 30 minutes (r=0.305, p=0.02). Moreover, plasma FSH levels were also positively correlated with age (r=0.438, p<0.001), LH (r=0.753, p<0.001), estradiol (r=0.595, p<0.001), testosterone (r=0.373, p<0.001), and inversely with the waist-to-hip circumference ratio (r=-0.360, p=0.001).

Subgroup analysis of children without obesity, categorized by pubertal status, revealed inverse correlations between FSH levels and BMI-SDS (r=-0.681, p<0.001), waist-to-hip circumference ratio (r=-0.726, p=0.001), and waist circumference-to-height ratio (r=-0.729, p=0.001) in prepubertal children. However, these relationships were not significant in pubertal children (data not shown). In pubertal children, FSH was positively correlated with S1PhOGTT (r=0.292, p=0.03) and S2PhOGTT (r=0.291, p=0.04), although no significant relationship was observed between FSH and these indices in prepubertal children (data not shown).

##### 3.4.1.3. Comparative analyses in children with obesity

Clinical and laboratory characteristics of children with obesity, categorized by pubertal status at their initial visit, are summarized in Table 2. Although prepubertal children presented with higher BMI-SDS, pubertal children had greater insulin secretory indices, including S1PhOGTT and S2PhOGTT, and higher IR, as indicated by HOMA-IR. While fasting glucose levels were not significantly different, glucose levels were lower during the OGTT in pubertal children from minute 30 to 180.

**Table 2.** Cross-sectional comparison of clinical and laboratory characteristics in prepubertal and pubertal children with obesity at their initial visit.

|  | Prepubertal<br>(n=165) | Pubertal<br>(n=357) | p-value* |
| --- | --- | --- | --- |
| Age (years) | 9.6 (8.3-10.9) | 13.1 (11.3-14.6) | **<0.001** |
| Sex (%) | 95 boys (58%)/70 girls (42%) | 247 boys (69%)/ 110 girls (31%) | **0.009** |
| BMI-SDS | 3.2 (2.8-3.6) | 2.9 (2.5-3.2) | **<0.001** |
| WC/height | 0.64 (0.60-0.67) | 0.63 (0.58-0.67) | **0.02** |
| WC/HC | 0.99 (0.95-1.02) | 0.96 (0.91-1.01) | **<0.001** |
| LH (IU/L) | 0.1 (0.1-0.1) | 2.5 (1.1-4.18) | **<0.001** |
| FSH (IU/L) | 0.89 (0.54-1.48) | 3.0 (2.0-4.4) | **<0.001** |
| Estradiol (pmol/L) | 40 (40-40) | 62.6 (40-110.1) | **<0.001** |
| Testosterone | 0.42 (0.31-0.42) | 1.31 (0.50-7.58) | **<0.001** |
| Fasting glucose (mmol/L) | 5.4 (5.2-5.7) | 5.3 (4.9-5.8) | 0.15 |
| OGTT glucose 5 min (mmol/L) | 5.8 (5.4-6.2) | 5.7 (5.3-6.2) | 0.19 |
| OGTT glucose 10 min (mmol/L) | 6.6 (6.1-7.1) | 6.5 (5.9-7.2) | 0.23 |
| OGTT glucose 15 min (mmol/L) | 7.6 (6.9-8.2) | 7.4 (6.7-8.1) | 0.15 |
| OGTT glucose 30 min (mmol/L) | 8.8 (8.0-9.8) | 8.5 (7.7-9.4) | **0.03** |
| OGTT glucose 60 min (mmol/L) | 7.7 (6.8-8.8) | 7.4 (6.4-8.5) | **0.04** |
| OGTT glucose 90 min (mmol/L) | 7.5 (6.5-8.4) | 7.0 (6.2-7.9) | **<0.001** |
| OGTT glucose 120 min (mmol/L) | 7.2 (6.7-8.1) | 7.1 (6.2-7.8) | **0.01** |
| OGTT glucose 150 min (mmol/L) | 7.0 (6.4-7.9) | 6.6 (5.8-7.4) | **0.003** |
| OGTT glucose 180 min (mmol/L) | 6.1 (5.1-6.8) | 5.4 (4.7-6.4) | **0.002** |
| Fasting insulin | 109 (66-153) | 128 (90-195) | **<0.001** |
| OGTT insulin 5 min | 188 (95-300) | 188 (129-292) | 0.19 |
| OGTT insulin 10 min | 403 (250-646) | 424 (271-639) | 0.54 |
| OGTT insulin 15 min | 653 (403-1014) | 695 (438-1070) | 0.31 |
| OGTT insulin 30 min | 927 (655-1412) | 1042 (722-1660) | **0.045** |
| OGTT insulin 60 min | 590 (424-1063) | 764 (455-1195) | **0.030** |
| OGTT insulin 90 min | 597 (427-986) | 625 (375-1007) | 0.69 |
| OGTT insulin 120 min | 667 (403-1042) | 636 (342-1070) | 0.69 |
| OGTT insulin 180 min | 323 (172-738) | 215 (101-479) | **0.004** |
| Delta insulin 0-30 min | 831 (527-1273) | 906 (604-1485) | 0.13 |
| Insulinogenic index | 2.11 (1.43-3.03) | 2.33 (1.58-3.50) | 0.056 |
| S1PhOGTT | 2183 (1622-3199) | 2558 (1796-3733) | **0.004** |
| S2PhOGTT | 548 (413-784) | 630 (455-911) | **0.004** |
| HOMA-IR | 3.60 (2.11-5.20) | 4.30 (2.82-6.6) | **<0.001** |

Abbreviations: BMI-SDS: Body mass index standard deviation score; FSH: Follicle-stimulating hormone; HC: Hip circumference; HOMA-IR: Homeostatic model assessment for insulin resistance; LH: Luteinizing hormone; min: Minutes; OGTT: Oral glucose tolerance test; S1PhOGTT: Stumvoll index 1; S1PhOGTT: Stumvoll index 2; WC: Waist circumference. All values are median and (Q1-Q3 interquartile ranges), except if otherwise stated. \*Bold text indicates a statistically significant difference with a p-value less than 0.05.

##### 3.4.1.4. Correlation and regression analyses in children with obesity

FSH levels showed a positive correlation with delta insulin (r=0.113, p=0.02), S1PhOGTT (r=0.139, p=0.003), S2PhOGTT (r=0.138, p=0.003) and HOMA-IR (r=0.141, p=0.002). Additionally, FSH levels were positively correlated with fasting insulin (r=0.150, p=0.001), and insulin values during the OGTT at 30 minutes (r=0.122, p=0.009) and 60 minutes (r=0.150, p=0.001). Interestingly, FSH was inversely correlated with glucose levels during the OGTT at 90 minutes (r=-0.119, p=0.01), 150 minutes (r=-0.310, p=0.002), and 180 minutes (r=-0.225, p<0.001). Blood FSH levels showed positive correlations with age (r=0.424, p<0.001), LH (r=0.729, p<0.001), estradiol (r=0.451, p<0.001), and testosterone (r=0.342, p<0.001). Additionally, FSH levels inversely correlated with the waist-to-hip circumference ratio (r=-0.213, p<0.001), BMI-SDS (r=-0.271, p<0.001), and the waist circumference-to-height ratio (r=-0.134, p=0.005).

In subgroup analyses of children with obesity, sorted by pubertal status, FSH levels were inversely correlated with BMI-SDS in prepubertal children (r=-0.371, p<0.001). However, this relationship was not observed in pubertal children (r=-0.044, p=0.4). In pubertal children, FSH was positively correlated with delta insulin (r=0.138, p=0.01), S1PhOGTT (r=0.131, p=0.02) and S2PhOGTT (r=0.128, p=0.02), although no significant relationship was observed between FSH and these indices in prepubertal children (data not shown). Additionally, positive correlations were noted between FSH and insulin values during the OGTT at 10 (r=0.135, p=0.02), 15 (r=0.126, p=0.03), 30 (r=0.134, p=0.02), 60 (r=0.158, p=0.004), and 90 minutes (r=0.121, p=0.03) in pubertal children. Furthermore, after controlling for BMI-SDS as a potential confounder for IR and insulinogenic indices, in children with obesity, the relationships between FSH and delta insulin (β=0.169, p=0.001), FSH and S1PhOGTT (β=0.148, p=0.004), and FSH and S2PhOGTT (β=0.149, p=0.004) were still significant, in pubertal children. Additionally, in these children, the relationships between FSH and HOMA-IR (β=0.103, p=0.038) along with FSH and IGI (β=0.117, p=0.027) were statistically significant after controlling for BMI-SDS.

#### 3.4.2. Follow-up visit, longitudinal data analyses

##### 3.4.2.1. Comparison of clinical and laboratory characteristics in children with obesity from prepubertal baseline to pubertal follow-up

In children with obesity, a comparison of clinical and laboratory characteristics from the prepubertal baseline to the follow-up visit during puberty was conducted (Table 3). The mean follow-up time was 3.2±1.5 years. Although a reduction in BMI-SDS was observed at follow-up in these pubertal children, insulin secretory indices including IGI, S1PhOGTT, and S2PhOGTT, as well as HOMA-IR, were increased. However, there were no changes in glucose concentrations, either at fasting or during the OGTT, between the baseline and follow-up visits.

**Table 3.** Longitudinal comparison of clinical and laboratory features in children with obesity from prepubertal baseline to follow-up visit during puberty (mean follow-up time:3.2±1.5 years).

|  | Prepubertal (First Visit)<br>(n=55) | Pubertal (Follow-up Visit)<br>(n=55) | p-value* |
| --- | --- | --- | --- |
| Age (years) | 9.5 (8.6-10.95) | 12.8 (11.8-13.7) | **<0.001** |
| Sex (%) | 32 boys (58%)/23 girls (42%) | 32 boys (58%)/ 23 girls (42%) | 1.0 |
| BMI-SDS | 3.2 (2.8-3.6) | 3.1 (2.65-3.2) | **0.006** |
| LH (IU/L) | 0.1 (0.1-0.1) | 3.25 (1.91-5.43) | **<0.001** |
| FSH (IU/L) | 0.89 (0.57-1.41) | 4.3 (2.7-5.63) | **<0.001** |
| Estradiol (pmol/L) | 40 (40-40) | 65 (40-116.3) | **<0.001** |
| Testosterone | 0.42 (0.31-0.42) | 1.31 (0.71-8.15) | **<0.001** |
| Fasting glucose (mmol/L) | 5.5 (5.2-5.7) | 5.7 (5.4-5.9) | 0.068 |
| OGTT glucose 5 min (mmol/L) | 5.8 (5.5-6.3) | 6.0 (5.6-6.4) | 0.68 |
| OGTT glucose 10 min (mmol/L) | 6.9 (6.4-7.4) | 6.6 (6.0-7.3) | 0.19 |
| OGTT glucose 15 min (mmol/L) | 7.7 (7.1-8.5) | 7.4 (7.0-8.3) | 0.15 |
| OGTT glucose 30 min (mmol/L) | 9.0 (8.2-10) | 8.7 (8.1-10.5) | 0.60 |
| OGTT glucose 60 min (mmol/L) | 7.8 (6.7-9.1) | 8.6 (7.5-10.7) | 0.054 |
| OGTT glucose 90 min (mmol/L) | 7.1 (6.4-9.2) | 8.0 (7.1-9.5) | 0.050 |
| OGTT glucose 120 min (mmol/L) | 7.4 (6.7-9.0) | 7.7 (6.7-8.8) | 0.11 |
| OGTT glucose 150 min (mmol/L) | 6.7 (6.2-7.6) | 7.1 (5.5-7.6) | 0.86 |
| OGTT glucose 180 min (mmol/L) | 6.2 (5.2-6.6) | 5.7 (4.9-7.3) | 0.87 |
| Fasting insulin | 117 (83-160) | 191 (117-302) | **<0.001** |
| OGTT insulin 5 min | 188 (106-326) | 320 (165-424) | **0.029** |
| OGTT insulin 10 min | 431 (300-762) | 445 (285-745) | 0.84 |
| OGTT insulin 15 min | 698 (455-1153) | 903 (375-1229) | 0.80 |
| OGTT insulin 30 min | 938 (715-1462) | 1368 (729-2195) | 0.061 |
| OGTT insulin 60 min | 653 (412-1089) | 1056 (667-2000) | **0.008** |
| OGTT insulin 90 min | 698 (429-945) | 1056 (691-1729) | **0.001** |
| OGTT insulin 120 min | 889 (451-1191) | 1083 (667-1847) | 0.093 |
| OGTT insulin 180 min | 517 (264-818) | 410 (236-1007) | 0.307 |
| Delta insulin 0-30 min | 853 (617-1327) | 1153 (590-1806) | 0.13 |
| Insulinogenic index | 2.11 (1.49-3.37) | 2.61 (1.6-5.3) | **0.026** |
| S1PhOGTT | 2268 (1691-3243) | 3105 (2262-5191) | **0.011** |
| S2PhOGTT | 565 (437-796) | 765 (577-1233) | **0.009** |
| HOMA-IR | 3.82 (2.33-5.63) | 7.03 (4.12-10.37) | **<0.001** |

Abbreviations: BMI-SDS: Body mass index standard deviation score; FSH: Follicle-stimulating hormone; HC: Hip circumference; HOMA-IR: Homeostatic model assessment for insulin resistance; LH: Luteinizing hormone; min: Minutes; OGTT: Oral glucose tolerance test; S1PhOGTT: Stumvoll index 1; S1PhOGTT: Stumvoll index 2; WC: Waist circumference. All values are median and (Q1-Q3 interquartile ranges), except if otherwise stated. \*Bold text indicates a statistically significant difference with a p-value less than 0.05.

##### 3.4.2.2. Correlation and regression analyses in children with obesity during pubertal follow-up

There were no significant correlations between FSH and BMI-SDS, FSH and age or FSH and HOMA-IR at the follow up visit during puberty (data not shown). However, plasma FSH levels were positively correlated with delta insulin (r=0.387, p=0.02) and IGI (r=0.323, p=0.045) at the follow-up. Additionally, FSH levels were positively correlated with insulin values during the OGTT at 5 minutes (r=0.329, p=0.04), 15 minutes (r=0.380, p=0.02), and 30 minutes (r=0.364, p=0.02). The associations between FSH levels and S1PhOGTT as well as S2PhOGTT did not achieve statistical significance with simple correlation analyses (data not shown). After controlling for BMI-SDS as a potential confounder, the relationships between FSH and delta insulin (β=0.491, p=0.008) and FSH and IGI (β=0.476, p=0.009) remained significant, and the relationships between FSH and S1PhOGTT (β=0.497, p=0.005), and FSH and S2PhOGTT (β=0.493, p=0.005) became significant.

## Discussion

Since the discovery of FSH in the early 20^th^ century, its central role in reproductive biology has been well documented^20^. Remarkably, even after approximately 100 years of research, the biology of FSH and its receptor FSHR remain an expansive field for exploration, underscoring the complexity and depth of the underlying mechanisms^20^. Our current study expands the traditional understanding of FSH’s role, revealing its direct influence on metabolic pathways through its effects on insulin secretion in the human pancreas. We demonstrated FSHR expression in human pancreatic beta-cells and showed that FSHR activation not only enhances GSIS but also induces a dose-dependent increase in cAMP production in isolated human pancreatic islets. Additionally, in line with our preliminary data from the ULSCO cohort^19^, findings in the present study from the extended Beta-JUDO cohort reinforce the view that FSH might be intricately involved in the modulation of insulin secretion, especially during puberty.

FSH, an evolutionarily well conserved hormone, has ancient origins, tracing back to species as early as jawed teleost fishes^47^. Such conservation suggests that FSH may have multiple, yet unrecognized, roles in mammalian physiology. Our findings of FSHR co-localization with insulin-producing beta-cells in human pancreatic islets are in line with prior discoveries of pancreatic FSHR expression in diverse species, including the Chinese alligator^48^, rat^24^ and mouse^49^. Additionally, the stimulatory effects of FSH on insulin secretion from human islets observed in the present study are supported by observations in rats^24^, female canines^50^ and mice^49^, showing direct effects of FSH on insulin secretion. Furthermore, a very recent study by Cheng et al.^49^ also reported FSHR expression in human pancreatic islets and showed direct regulatory effects of FSH on insulin secretion. Their findings, which are in line with our own previous^25^ and present findings, indicated that insulin secretion is enhanced by FSH at high glucose concentrations. However, unlike our results, the study by Cheng et al.^49^ reported that the relationship between FSH concentration and its promotion of GSIS exhibited a bell curve pattern. While FSH levels up to 10 IU/L augmented GSIS dose dependently, concentrations at 100 IU/L attenuated GSIS^49^. The authors also observed that treatment with FSH concentrations below 10 IU/L markedly increased intracellular cAMP levels in a concentration-dependent manner, whereas high concentration of FSH at 100 IU/L decreased the intracellular cAMP levels^49^. These results contrast with our observations of a dose-dependent increase in cAMP in response to FSH under stimulatory glucose conditions. These discrepancies could be attributed to the different experimental methods used. Additionally, while Cheng et al.^49^ used human pancreatic islets solely for expression analysis and performed functional experiments using either mouse pancreatic islets or the mouse insulinoma cell line MIN6, we used human pancreatic islets for all experimental investigations.

The observed potentiation of GSIS by FSH in our study suggests a possible involvement in both normal physiological regulation and pathological states of metabolic processes. Aligning with previous research^10^, our clinical data showed that blood FSH levels during puberty fall within the stimulatory range for insulin secretion as demonstrated *in vitro*. These findings, also supported by the recent article by Cheng et al.^49^, might imply that the increased levels of FSH associated with puberty may play a role in the emergence of hyperinsulinemia and IR during this critical period. Additionally, the detection of FSHR in the human adipose tissues^22^, and the noted effects of interventions that block FSH action in reducing adiposity in mice models^23^, hint at a broader metabolic role for FSH. Consequently, the direct effects of FSH on both the increase in insulin secretion and adipose tissue IR may establish a feedback loop that contributes to the development and progression of these metabolic processes.

From an evolutionary perspective, metabolism, growth, and reproduction are tightly interconnected^51^. Fertility is typically reached after completing puberty and growth, and it is deeply influenced by the individual’s metabolic status. The complex interplay between these processes is believed to arise from shared regulatory networks and signaling pathways, which are not yet fully understood^51^. FSH with its direct effects on gonads and insulin secretion could be one of the key elements in this interplay. Puberty is marked by rapid weight gain and growth spurt^11^, aligning with the beginning of reproductive capacity and significant metabolic changes to accommodate increased energy demands^52^. In agreement with our findings, it is well established that both fasting and prandial insulin concentrations rise during puberty in children, regardless of whether they have normal weight or obesity^12, 14, 15^. This puberty-related surge in insulin has been linked to the rapid weight gain typically observed during this developmental phase^12^. Although weight gain is physiological and expected during puberty, the metabolic changes that naturally occur may put adolescents at increased risk for excessive weight gain^13^. Furthermore, evidence suggests that puberty is an important risk factor for transitioning from metabolically healthy to unhealthy obesity^14, 19^. Our analyses, which showed associations of FSH with insulin secretion indices independent of BMI in pubertal children, provide further evidence suggesting that elevated FSH levels during puberty might contribute to puberty-related insulin hypersecretion and IR. In susceptible individuals, these elevated insulin levels may also lead to the development of metabolic derangements during this critical period^19^. Increased insulin levels may also influence other physiological changes during puberty. Elevated gonadotropins boost sex steroid production, a key process in pubertal development^1^. FSH may also contribute to this progression through its direct effect on insulin hypersecretion, which can lower sex hormone-binding globulin levels and consequently increase the availability of free sex hormones^53^.

Nonetheless, our study has limitations. The low number of prepubertal children without obesity and the lack of longitudinal data for these children limit the generalizability of our findings across different body weight categories. Moreover, the complex interplay between FSH and other fluctuating hormones during puberty, such as LH, sex steroids, GH, and IGF-1, necessitates cautious interpretation of the isolated effects of FSH. An additional limitation is the age disparity between our pediatric clinical participants and the primarily adult donors of the pancreatic islet cells used in our experiments. Additionally, we did not investigate FSHR expression in other pancreatic endocrine and exocrine cells or its effects on other hormone productions.

In conclusion, we found that FSH directly influences pancreatic beta-cell function and likely contributes to the modulation of insulin secretion during puberty. This study expands the conventional understanding of FSH to include significant metabolic functions, particularly during this critical period. The results also set a clear path for future studies aiming to further decipher FSH’s metabolic role. Comprehensive longitudinal research, encompassing a wider range of age and weight categories, is essential to detail FSH’s specific effects on metabolic alterations. Additionally, investigating the complex interplay of FSH with other critical hormones is crucial to comprehending their combined effects on metabolic regulation.

## Acknowledgements

We thank the participants of the Beta-cell function in Juvenile Diabetes and Obesity (Beta-JUDO) cohort and their families. Additionally, we sincerely appreciate the contributions of the staff at the childhood obesity clinics in Uppsala and Salzburg.

This study was funded by the European Commission’s Seventh Framework Programme (FP7/2012–2018) project Beta-Judo (grant number 279153), Swedish Diabetes Association (grant number DIA 2016–146), Swedish Foundation for Strategic Research (grant number CMP22-0014), Sweden’s Innovation Agency Vinnova (grant number 2020-02417), Family Ernfors Foundation (grant number 160504), Swedish Research Council (Dnr 2016–01040), Excellence of Diabetes Research in Sweden (EXODIAB), Uppsala-Örebro Regional Research Council (RFR 158161, RFR 233041, RFR 309901), Gillbergska Foundation (Uppsala, Sweden), the Swedish Society for Diabetology, and the Selander Foundation, Swedish Research Council (2019-01456), Novo-Nordisk foundation (NNF19OC0055275), Swedish Diabetes Association (DIA2023-781).

## Author contributions

B.K.A., P.B., A.C. and O.I.H. conceived and designed the *in vitro* experiments. P.B., A.F. and D.W. conceived and designed the in vivo experiments. P.B., O.I.H., A.F. and D.W. provided experimental infrastructure. B.K.A., C.I.N., A.C., Q.W., S.Y.C. and R.S. performed the *in vitro* experiments. C.I.N. and O.I.H. performed and analyzed FSH-mediated signaling network experiments. R.S., K.M., I.C., H.M. performed the *in vivo* experiments. B.K.A. and C.I.N. wrote the manuscript. All authors participated in interpreting the results and revising the manuscript.

